# SF3B1ness score: screening *SF3B1* mutation status from over 60,000 transcriptomes based on a machine learning approach

**DOI:** 10.1101/572834

**Authors:** Yuichi Shiraishi, Kenichi Chiba, Ai Okada

## Abstract

In precision oncology, genomic evidence is used to determine the optimal treatment for each patient. However, identification of somatic mutations from genome sequencing data is often technically difficult and functional significance of somatic mutations is inconclusive in many cases. In this paper, to seek for an alternative approach, we tackle the problem of predicting functional mutations from transcriptome sequencing data. Focusing on *SF3B1*, a key splicing factor gene, we develop SF3B1ness score for classifying functional mutation status using a combination of Naive Bayes classifier and zero-inflated beta-binomial modeling (R package is available at (https://github.com/friend1WS/SF3B1ness). Using 8,992 TCGA exome and RNA sequencing data for evaluation, we show that the classifier based on SF3B1ness score is able to (1) attain very high precision (>93%) and sensitivity (>95%), (2) rescue several somatic mutations not identified by exome sequence analysis especially due to low variant allele frequencies, and (3) successfully measure functional importance for somatic mutation whose significance has been unknown. Furthermore, to demonstrate that the SF3B1ness score is highly robust and can be extensible to the cohorts outside training data, we performed a functional *SF3B1* mutation screening on 51,577 additional transcriptome sequencing data. We have detected 135 samples with putative *SF3B1* functional mutations including those that are rarely registered in the somatic mutation database (e.g., G664C, L747W, and R775G). Moreover, we could identify two cases with *SF3B1* mutations from normal tissues, implying that SF3B1ness score can be used for detecting clonal hematopoiesis.

## Introduction

Advances in high throughput sequencing technology are bringing about revolutions in medicine. By sequencing each patient’s cancer genomes, clinicians can discover appropriate treatment specifically targeted to the detected somatic variants. On the other hand, somatic mutations profiled via genome sequencing is usually far from perfect. It is still difficult to identify somatic mutations with complex forms [25, 27] and low variant allele frequencies [21]. In addition, although a vast number of cancer genome studies revealed a huge number of novel cancer-driving genes, the functional significance of variant for each position is still uninvestigated even for established cancer genes such as *BRCA1*/*2* [2] and *EGFR* [11].

Using RNA sequencing together with genome sequencing often help complementing somatic mutation discovery and measuring the effect of a genomic mutation on transcriptomes such as expression and splicing changes. Here, we would tackle the problem of classifying “functional” mutation status based on transcriptome using statistical machine learning approaches.

In this paper, we would like to focus on *SF3B1* gene, which encodes a core component of the RNA splicing machinery. *SF3B1* is recurrently mutated in blood cancers [29] and other solid tumors [7, 17]. The somatic mutations periodically cluster to positions within C-terminal HEAT repeat domains (HDs) with several major hotspots including p.R625, p.K666, p.K700 [20]. Although previous studies have clarified the functional significance for a few prominent positions [1], and the other sites that are less frequently but recurrently mutated are little investigated.

Figure 1A illustrates the outline of this study. We first develop a novel statistical approach that can effectively capture the characteristics of splicing changes induced by *SF3B1* mutations using zero-inflated beta-binomial distribution naive Bayes classifier. Then, applying the proposed method to TCGA transcriptome sequencing data, we fit and evaluate the model on samples across 31 cancer types. Through that process, we provide a novel measure, SF3B1ness score, that quantifies transcriptional abnormalities caused by *SF3B1* functional mutations in a unified way. Then, through the use of SF3B1ness score on about 60,000 publicly available transcriptome sequencing data, we perform large scale screening of *SF3B1* functional mutations.

**Figure 1.**
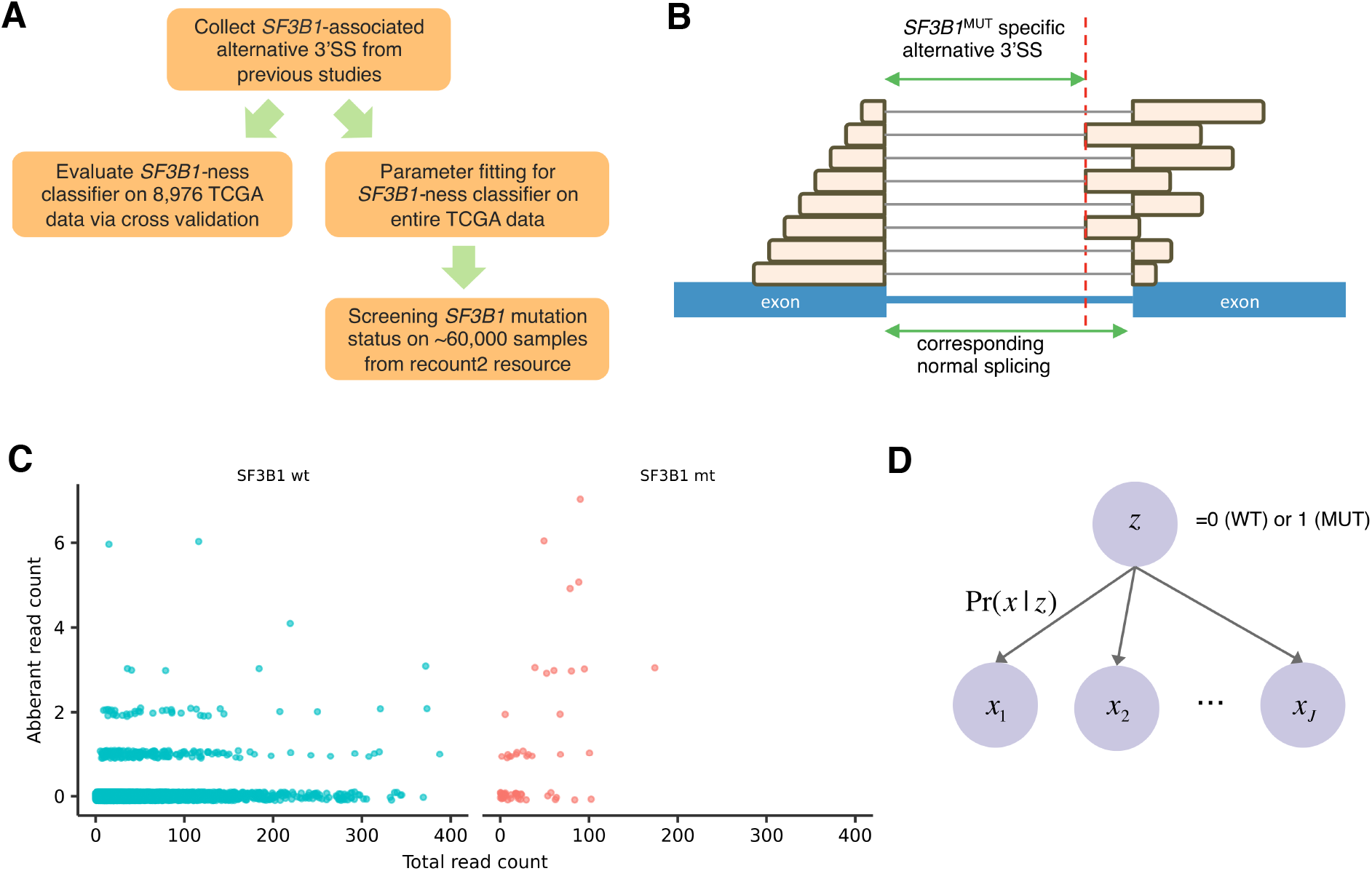
Overview of the study. **A**, The process of evaluating SF3B1ness classifier and screening functional *SF3B1* mutations. **B**, Illustration of *SF3B1*^MUT^ associated alternative 3’SS and its corresponding normal splicing identified by split-aligned short reads. **C**, An example of supporting read counts of alternative 3’SS (chr1:100477090-100480840) and its corresponding normal splicing (chr1:100477090-100480857). Each point shows the counts of an individual. **D**, The generative model of assumed in this paper. The counts of *SF3B1*^MUT^ associated alternative 3’SS supporting reads are conditionally independent given the *SF3B1* mutation status.

## Result

### Method overview

It has been known that functional *SF3B1* mutations generate hundreds to thousands of genome-wide aberrant splicing events and these are mostly alternative 3’SS (splice site), where one end of splicing junction is at the annotated splicing donor site and the other end is distantly located from the annotated splicing acceptor site (see Figure 1B). Therefore, we first collected the set of these *SF3B1*^MUT^ associated alternative 3’SS events from two previous studies [1, 4], and use the quantification of these events (the numbers of short sequence reads spanning the target splicing junction [14, 23]) as the features for the classification problem. In addition, the count of the corresponding normal splicing event for each alternative 3’SS is also used for measuring the ratio of alternative 3’SS (see Material Method for detail). In the end, we could extract 710 pairs of *SF3B1*^MUT^ associated alternative 3’SS and corresponding normal splicing.

Calculations of splicing junction events require downloading and alignment of raw sequencing data and thus needs considerable storage and computational cost. However, there is an excellent resource, the recount2 [3], which provides already well-processed transcriptome data including splicing junction counts for ≥ 70,000 samples from ≥ 2,000 different studies including The Cancer Genome Atlas (TCGA) and The Genotype-Tissue Expression (GTEx) project. Therefore, to quantify *SF3B1*^MUT^ associated alternative 3’SS, we just downloaded the splicing junction data from the recount2 for each study and extracted those matching the *SF3B1*^MUT^ associated alternative 3’SS and their corresponding normal splicing.

For typical *SF3B1*^MUT^ associated alternative 3’SS loci, aberrant read counts are zeros in most *SF3B1*^WT^ samples, whereas some samples have a decent amount of aberrant read counts (see Figure 1C). Even though the ratios of aberrant read counts significantly increase for many *SF3B1*^MUT^ samples, still a certain number of samples have zero counts. In fact, these excess zero-count situations cannot be not effectively captured by common probabilistic distributions such as binomial distribution, and zero-inflated models (mixtures of probability mass at zero and other distributions) have been used in many biological data analysis such as single-cell transcriptome [10, 19] and microbiome [8, 18, 26]. In this paper, we assume that the counts of *SF3B1*^MUT^ associated alternative 3’SS and corresponding normal splicing are generated by zero-inflated beta-binomial distribution (see Material Method for detail).

For constructing the classifier on *SF3B1* mutation status from alternative 3’SS count data, we adopt naive Bayes classifier (see Figure 1D), The detailed procedure is as follows (see Material Method for detail):

1. We divide the samples into *SF3B1*^WT^ and *SF3B1*^MUT^ groups. Then, for each group and each *SF3B1*^MUT^ associated alternative 3’SS, we estimate the parameters of zero-inflated beta-binomial model.
2. Assuming the conditional independence among the *SF3B1*^MUT^ associated alternative 3’SS events, we calculate the posterior probabilities of *SF3B1*^WT^ and *SF3B1*^MUT^ based on the parameters estimated above
3. By taking the difference of the log probabilities of *SF3B1*^MUT^, which we define as “SF3B1ness” score, we predict the *SF3B1* mutation status (positive if and only if the SF3B1ness score is above a threshold).

### Application to TCGA data

To evaluate the effectiveness of SF3B1ness classifier, we applied the proposed approach to TCGA data set. We selected the set of 8,992 primary cancer samples from 31 types and determined the somatic mutation status for *SF3B1* as described in the previous study [23]. In this section, referring to past literature, we consider the *SF3B1* hotspot mutations as those occurring at E622, R625, N626, H662, K666, K700, G740, K741, G742. Then, there are 63 samples with SF3B1 hotspot mutations in total. The cancer types with frequent *SF3B1* hotspot mutations are breast invasive carcinoma (BRCA, 14 / 826), skin cutaneous melanoma (SKCM, 8 / 469), and uveal melanoma (UVM, 17 / 80). Besides, there are other cancer types with *SF3B1* hotspot mutations albeit the frequencies is low including bladder urothelial carcinoma, kidney renal clear cell carcinoma, lung adenocarcinoma, prostate adenocarcinoma, and so on.

We divided 8,992 samples into two datasets and used one dataset (training data) for fitting the parameters of SF3B1ness classifier and obtained the SF3B1ness score for each sample in the other data set (test data). Then, we performed the same procedure exchanging the training and test data. When training the parameters, samples with well known *SF3B1* hotspot are categorized as positive cases (*z_i_* = 1), and those without any *SF3B1* mutations are categorized as negative cases (*z_i_* = 0). Note that samples with non-hotspot *SF3B1* mutations are removed in the training phases because these biological functions remain to be investigated and only used in test phases. Importantly, we used samples from all the 31 cancer types together without any distinction. Although the amount of expression of each *SF3B1*^MUT^ associated alternative 3’SS varied according to cancer types, we believe that our model is sufficiently robust to these variations. The key to this robustness is that we take not only the absolute count of *SF3B1*^MUT^ associated alternative 3’SS but also the fraction to its normal splicing into account via beta-binomial distribution. Furthermore, the adoption of zero-inflated components makes the probabilistic model tolerant of other disturbing factors such as outliers and mislabeling.

First, the mutation status could be very accurately predicted by the proposed method. Most of the samples with high SF3B1ness score actually had *SF3B1* hotspot mutations (see Figure 2A and S1 Table). Among 63 samples with known *SF3B1* hotspot mutations, 60 samples had positive SF3B1ness score, indicating that the sensitivity of *SF3B1* score is more than 95%.

**Figure 2.**
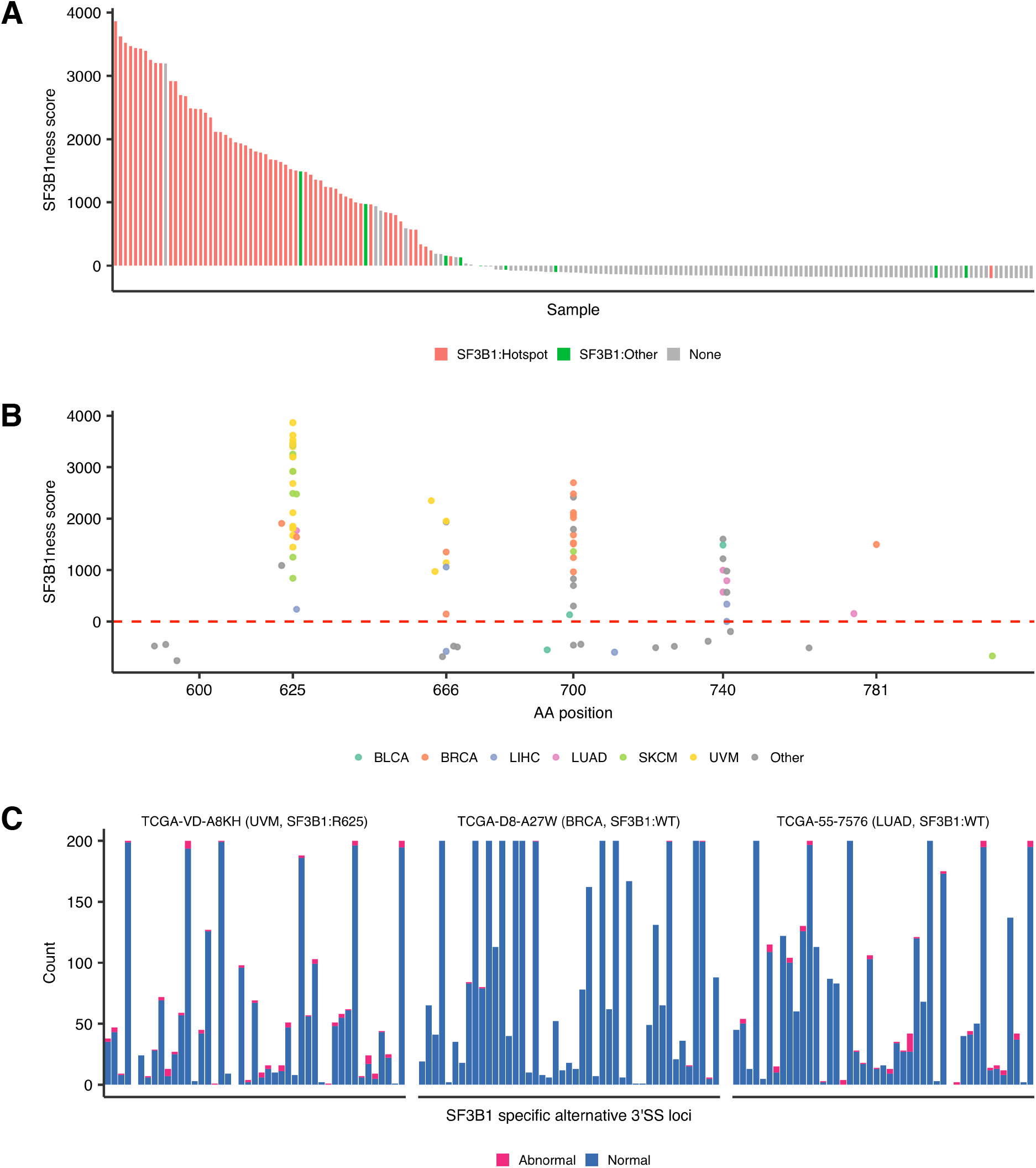
The overview of SF3B1ness score evaluation on TCGA data. **A**, The relationships between SF3B1ness scores and *SF3B1* mutation status. Each bar shows each individual from TCGA dataset, and the color shows *SF3B1* mutation status (hotspot are defined in the above). **B**, The relationships between SF3B1ness scores and amino-acids positions for samples with *SF3B1* mutations. The color shows the cancer types. **C**, The counts of *SF3B1*^MUT^ associated alternative 3’SS and its corresponding normal splicing junction for several characteristic samples. Each bar shows the loci of *SF3B1*^MUT^ associated alternative 3’SS. For loci with ≥ 200 supporting read counts in total, they are down-sampled to 200.

There were 13 samples with positive *SF3B1* score even without hotspot *SF3B1* mutations (see Table 1). Those samples are categorized into the following three classes.

1. Four samples had non-hotspot *SF3B1* mutations (such as T663P, Q699E, R775L, and D781E). Although many of them have been comparably rarely detected in cancer genome studies, decent amounts of SF3B1ness score suggest that they have similar functions to hotspot mutations on genome-wide splicing aberrations.
2. For other four samples, after manually investigating the exome and RNA sequencing data, we could detect a number of short reads supporting *SF3B1* hotspot somatic mutation even though the variant allele frequencies are very low (1% − 10%). Therefore, even with low variant allele frequencies, the function of *SF3B1* somatic mutation can be detectable.
3. For the remaining five samples, we could not identify any *SF3B1* putative functional mutations after extensive manual investigations. Still, the profiles of alternative 3’SS counts for these samples are very similar to those with hotspot *SF3B1* mutations (see Figure 2C), implying that there may be other genomic mutations on the same pathway with *SF3B1* concealed in these samples.

**Table 1.** List of samples with high SF3B1ness score and without hotspot *SF3B1* mutations.

| Sample name | Cancer type | Score | Remark |
| --- | --- | --- | --- |
| TCGA-55-7576 | LUAD | 3193.496 | No *SF3B1* mutation is identified after manual investigations. (low variant allele frequency). |
| TCGA-E2-A10F | BRCA | 1495.878 | D781E mutation is identified by usual exome data analysis. |
| TCGA-V4-A9EC | UVM | 972.768 | T663P mutation is identified by usual exome data analysis. |
| TCGA-3H-AB3K | MESO | 935.228 | No *SF3B1* mutation is identified after manual investigations. |
| TCGA-XM-A8RI | THYM | 866.908 | K700E mutation is identified after manual investigations. (low variant allele frequency). |
| TCGA-05-4432 | LUAD | 586.704 | No *SF3B1* mutation is identified after manual investigations. |
| TCGA-75-5125 | LUAD | 190.175 | K700E mutation is identified after manual investigations. (low variant allele frequency). |
| TCGA-ZP-A9CV | LIHC | 186.524 | No *SF3B1* mutation is identified after manual investigations. |
| TCGA-NJ-A55O | LUAD | 153.664 | R775L mutation is identified by usual exome data analysis. |
| TCGA-NJ-A4YQ | LUAD | 136.287 | No *SF3B1* mutation is identified after manual investigations. |
| TCGA-UY-A78N | BLCA | 130.962 | Q699E mutation is identified by usual exome data analysis. |
| TCGA-EY-A2OM | UCEC | 38.213 | No *SF3B1* mutation is identified after manual investigations. |
| TCGA-WC-A87W | UVM | 21.388 | R625H mutation is identified after manual investigations. (low variant allele frequency). |

The *SF3B1* somatic mutations linked to high SF3B1ness scores are in the proximity of known hotspots (see Figure 2B). Although this is anticipated by the marked concentration of *SF3B1* mutations and oncogenic functions thereof. Still, SF3B1ness score discerned the amino acid positions into those with high (e.g., T663P, Q699E) from those with low (e.g., I665F, V668I, G693) scores even among the proximity of hotspots.

Collectively, SF3B1ness score is helpful for measuring the effects of rare *SF3B1* mutations as well as saving mutations misidentified because of some technical problems such as low variant allele frequencies. Assuming that samples belonging to the first two classes are true positives, SF3B1ness score has fairly high precision (93.15%) on the functional *SF3B1* mutation status prediction.

### Screening of *SF3B1* functional mutations from large scale transcriptome resource

To evaluate the ability of SF3B1ness score on dataset not seen during training, we performed further large scale screening of functional *SF3B1* mutation from in total of 51,577 transcriptome data in the recount2 resource, where we removed data with small sizes (with ≤ 100M bases). We first calculated SF3B1ness score just by downloading splicing junction data (which is far more small in size than raw sequencing data). Then, for the samples with high SF3B1ness scores, we downloaded their raw sequence data, aligned to the reference genome and checked the *SF3B1* mutation status.

First, we identified 154 samples with ≥ 50.0 SF3B1ness scores. Of those, we could identify 140 samples having *SF3B1* mutations registered in the COSMIC database within C-terminal HEAT repeat domains (residues 604–801). When narrowing down the 124 samples with ≥ 300.0SF3B1ness scores, then 123 samples (123 / 124 = 0.992) had COSMIC registered *SF3B1* mutations, indicating that the SF3B1ness score is highly robust and can be utilized to a broad type of transcriptome data without caring tissue or cancer types (see Figure 3A and S2 Table).

**Figure 3.**
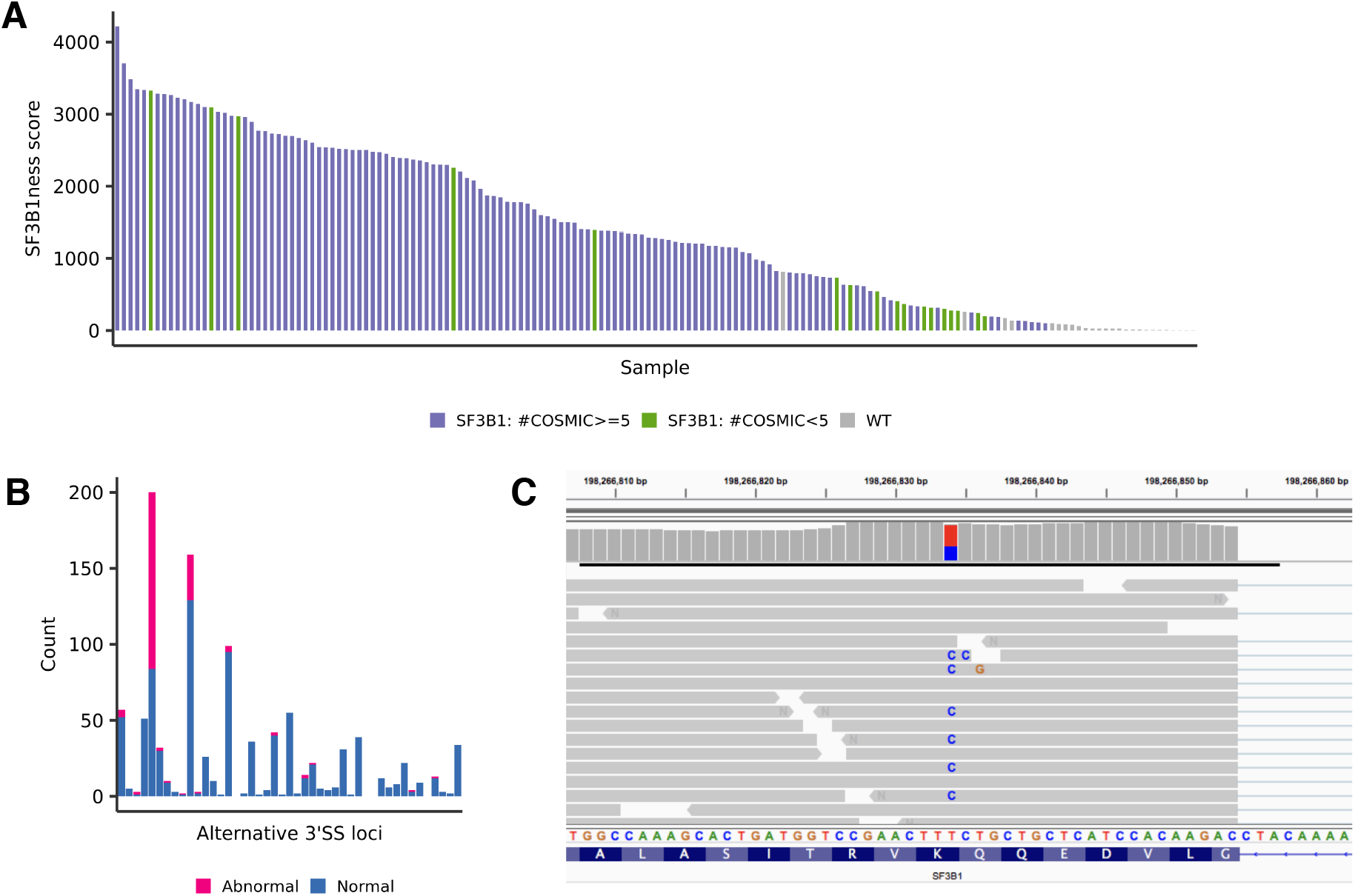
Overview of SF3B1 functional screening using the recount2. **A**, The relationships between SF3B1ness scores and *SF3B1* mutation status. Each bar shows each individual from recount2 dataset, and the color shows *SF3B1* mutation status (COSMIC release v87 is used for the registered number of *SF3B1* mutations). **B, C**, The counts of *SF3B1*_MUT_ associated alternative 3’SS and its corresponding normal splicing junction (B) and the alignment view around *SF3B1* K700E for the sample having high SF3B1ness score (SRR1374647) in GTEx study (C).

Many of the *SF3B1* mutations found in high SF3B1ness score samples are hotspot mutations (a large number of records registered in COSMIC database). However, there are several substitutions (R625G, G664C, K741N, L747W, and R775G) with few records in COSMIC, suggesting the functional importance of these rare mutations and awaiting for further biological experiments.

Most samples with high SF3B1ness scores are found in cancer types where *SF3B1* mutations are known to be prevalent, including AML (acute myeloid leukemia), CLL (chronic lymphocytic leukemia), MDS (myelodysplastic syndrome), breast cancer, melanoma, and uveal melanoma. However, we could also find *SF3B1* mutations from two normal samples; one is from a whole blood sample (SRR1374647, GTEX-ZF28-0005-SM-4WKH3) from GTEx study [15] (see Figure 3B,C). The other is from a bone sample from an old woman (SRR2305491) [6]. These facts indicate that *SF3B1* mutations induce splicing alterations in even normal tissues and SF3B1ness score may be helpful for efficient detection of clonal hematopoiesis from individuals without apparent hematological malignancies.

## Discussion

In this paper, we could constitute the classifier, SF3B1ness score, to accurately identify *SF3B1* functional mutation status. Most classifiers for predicting some pathway activation are trained on specific cancer types or tissues. However, the SF3B1ness score developed in this paper is highly robust so that users can easily apply it to their own data without caring the properties target transcriptome sequencing data. This will facilitate the application of SF3B1ness score for screening *SF3B1* mutation status from large scale transcriptome data. The robustness is partly because the target loci of splicing changes by *SF3B1* mutations are mostly common across cancer types and tissues. However, at the same time, the delicately designed probabilistic model (e.g., adopting zero-inflation components) significantly contribute to this robustness.

There have been active researches on inhibiting SF3B1 as a therapeutic target [13], and several studies demonstrated that mutations within *SF3B1* impact on the sensitivities to SF3B1 inhibitors [16, 28], suggesting that the SF3B1ness proposed in this study may be helpful to evaluate the activity of aberrant splicing, and future precision medicine.

One may argue that directly investigating *SF3B1* mutation by aligning transcriptome sequencing data and checking the variant allele frequencies of known hotspots is much easier and more straightforward. For that opinion, we would like to insist that automatically detecting somatic mutation is still not trivial especially for those with low variant allele frequencies and our approach can give another way to confirm and rescue *SF3B1* functional mutations. In addition, since raw sequencing data usually includes information that is sufficient to identify individuals and we need to follow ethically appropriate procedures (which is often time-consuming) to access and manage raw sequencing data. On the other hand, several consortium and resource distribute already pre-processed splicing junction data [3, 9, 12, 15], where our approach offers perfectly simple and accurate way for somatic mutation screening.

There are other splicing factor genes such as *U2AF1, SRSF2*, and several studies show that *SETD2*, a histone methyltransferase, is also related to aberrant splicing [24]. In fact, the characteristics of splicing changes in these genes are largely different. For *SF3B1* mutation, completely novel splicing junctions are generated, whereas just the ratios of already splicing junctions change by the mutation in other genes. Therefore, to be able to constitute classifier for other genes related to splicing, the model developed in this paper need to be significantly refined.

## Materials and Methods

### Collection of *SF3B1*^MUT^ associated alternative 3’SS

Through a large number of studies [1, 4, 5, 22], the characteristics of *SF3B1*^MUT^ abnormal splicing are known as follows:

- These aberrant splicing events are mostly alternative 3’SS.
- Also, the new acceptor sites of *SF3B1*^MUT^ associated alternative 3’SS events are typically located within 18 to 50bp downstream from the authentic acceptor sites.

We have compiled *SF3B1*^MUT^ specific abnormal splicing junctions from two previous studies. The first set is 895 splicing junctions identified by comparing 35 *SF3B1*^MUT^ and 50 *SF3B1*^WT^ samples from chronic lymphocytic leukemia, breast cancer, skin melanoma and uveal melanoma [4]. The second set is 1,124 spicing junctions identified by comparing 16 *SF3B1*^MUT^ and 56 *SF3B1*^WT^ samples from uveal melanoma [1]. We gathered the splicing junctions from the two studies and removed splicing junctions matching with annotated introns defined by Gencode Version 19 (http://hgdownload.cse.ucsc.edu/goldenPath/hg19/database/wgEncodeGencodeBasicV19.txt.gz). Furthermore, we request that one end of splicing junction is at the annotated splicing donor site and the other is located within 50bp upstream of canonical splicing acceptor sites. Then, the remaining the coordinates of splicing junctions are converted to GRCh38 based positions.

### A generative model of splicing junction counts via zero-inflated beta-binomial distribution

Let *x_i,j_, y_i,j_* denote the counts of *j*-th *SF3B1*^MUT^ associated alternative 3’SS and the corresponding normal splicing for the *i*-th sample (*i* = 1, 2, ⋯, *I, j* = 1, 2, ⋯, *J*). Setting *n_i,j_* = *x_i,j_* + *y_i,j_*, then the probability function of *x_i,j_* is

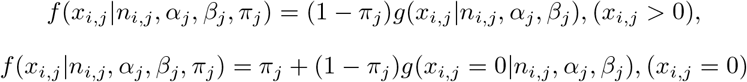

where *g* is the beta-binomial density function,

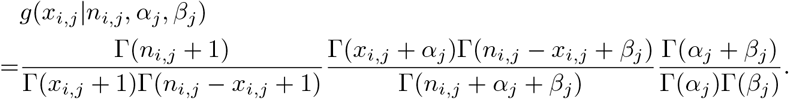

The parameters *α_j_, β_j_, π_j_* are estimated by numerically maximizing the log-likelihood,

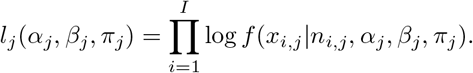

### A classification model for *SF3B1* mutation status using naive Bayes model

Suppose *z_i_* ∈ {0, 1} is the *SF3B1* mutation status for the *i*-th sample (0: *SF3B1*^WT^, 1: SF3B1^MUT^). First, for each *SF3B1*^MUT^ associated alternative 3’SS, we estimate the parameters of zero-inflated beta-binomial distribution for *SF3B1*^WT^ and *SF3B1*^MUT^ groups. Let 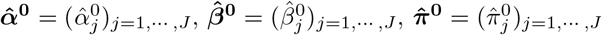 denote the parameters estimated for *SF3B1*^WT^ groups and 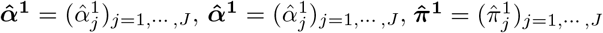 the parameters for *SF3B1*^MUT^ groups, respectively.

Then, by applying Bayes’ theorem, the conditional probabilities are

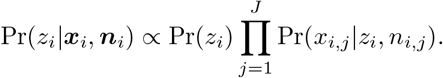

Therefore,

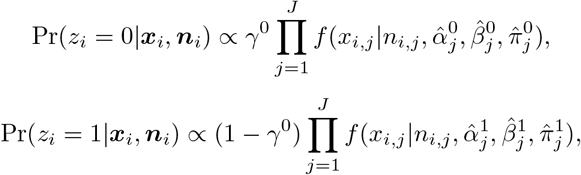

where *γ*^0^ is the parameter corresponding to Pr(*z_i_* = 0) (in this paper, we adopt non-informative value *γ*^0^ = 1/2). Finally, for each new sample, we evaluate the logarithm of the ratio of conditional probabilities (SF3B1ness score)

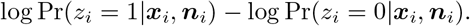

## Supporting Information

**S1 Table**.

**The SF3B1ness scores and mutation status for each TCGA samples**.

**S2 Table**.

**The SF3B1ness scores and mutation status for each recount2 samples with high SF3B1ness scores**. In the Mutation_Info columns, the amino-acids changes, the numbers of registered mutations in COSMIC database and variant allele frequencies are concatenated by commas. Also, if there are multiple somatic *SF3B1* mutations, these strings are linked together by semi-colons.

## Acknowledgments

The authors initiated this research project inspired by discussions with Dr. Claus Thorn Ekstrøm and Dr. Xiang Zhou through “No disease is an island” project, which is financed by the Danish Agency for Science and Higher Education. We would like to thank them.

